# Human cytomegalovirus induces neuronal gene expression for viral maturation

**DOI:** 10.1101/2024.06.13.598910

**Authors:** Laurel E Kelnhofer-Millevolte, Julian R Smith, Daniel H Nguyen, Lea S Wilson, Hannah C Lewis, Edward A Arnold, Mia R Brinkley, Adam P Geballe, Srinivas Ramachandran, Daphne C Avgousti

## Abstract

Viral invasion of the host cell causes some of the most dramatic changes in biology. Human cytomegalovirus (HCMV) extensively remodels host cells, altering nuclear shape and generating a cytoplasmic viral-induced assembly compartment (vIAC). How these striking morphology changes take place in the context of host gene regulation is still emerging. Here, we discovered that histone variant macroH2A1 is essential for producing infectious progeny. Because virion maturation and cellular remodeling are closely linked processes, we investigated structural changes in the host cell upon HCMV infection. We discovered that macroH2A1 is necessary for HCMV-induced reorganization of the host nucleus, cytoskeleton, and endoplasmic reticulum. Furthermore, using RNA-seq we found that while all viral genes were highly expressed in the absence of macroH2A1, many HCMV-induced host genes were not. Remarkably, hundreds of these HCMV-induced macroH2A1-dependent host genes are associated with neuronal synapse formation and vesicle trafficking. Knock-down of these HCMV-induced neuronal genes during infection resulted in malformed vIACs and smaller plaques, establishing their importance to HCMV infection. Together, our findings demonstrate that HCMV manipulates host gene expression by hijacking a dormant neuronal secretory pathway for efficient virion maturation.

## Introduction

Human Cytomegalovirus (HCMV) is a ubiquitous herpesvirus with seropositivity ranging from 65 to 100%^1,2^. In immunocompetent individuals, HCMV is typically asymptomatic or results in mild cold-like symptoms^3^. In contrast, congenital HCMV infection is one of the leading causes of infectious birth defects affecting about 1 in every 200 live births^4^. Additionally, cytomegalovirus is a chief cause of morbidity and mortality in both solid organ and stem cell transplant recipients^5^ and individuals with poorly controlled HIV^6^.

HCMV infection of the host cell induces large scale cellular remodeling^7^. The host nucleus forms a characteristic kidney-bean shape while host chromatin becomes polarized to one side^8^ and large-scale cytoskeletal rearrangement causes the nucleus to spin^8,9^. Simultaneously, the Golgi, endosome, and other cellular membranes are reorganized during HCMV infection to form the viral-induced assembly compartment (vIAC)^10^. Viral replication occurs in the nucleus, after which progeny capsids exit from the nucleus and pass through the vIAC for tegumentation and final maturation^11–13^. Importantly, unlike other lytic herpesvirus infections, HCMV does not shut down host transcription^14^, though how host gene expression is altered by HCMV remains actively under investigation.

Cellular remodeling is closely controlled by host gene expression, which in turn is regulated by histone modifications and histone variants^15–17^. MacroH2A1 is a histone variant that can replace the core histone H2A. MacroH2A1 was initially discovered on the inactive X chromosome associated with transcriptional repression^18,19^. In contrast, macroH2A1 is also required for gene activation in several contexts including serum starvation response^19^, smooth muscle differentiation^20^, and neuronal differentiation^21^. We previously showed how macroH2A1-dependent heterochromatin is critical for herpes simplex (HSV-1) egress from the nuclear compartment^22^. Therefore, we hypothesized that macroH2A1 may also function in cytomegalovirus infection. Here, we demonstrate that HCMV predominantly upregulates hundreds of exclusively neuronal genes in a macroH2A1-dependent manner. This upregulation of neuronal genes is essential for efficient HCMV infectious progeny production and defines a new mechanism wherein HCMV has evolved to control host gene expression to promote progeny maturation and viral spread.

## Results

### MacroH2A1 is required for production of infectious HCMV progeny

Due to the importance of macroH2A1 in lytic HSV-1 infection^22^ and the observation that macroH2A1 mRNA levels increase during lytic HCMV infection^23^, we hypothesized that macroH2A1 is also necessary for efficient HCMV lytic infection. To investigate this hypothesis, we infected wild-type human foreskin fibroblast cells (WT HFF-T) and our established macroH2A1 CRISPR knock-out cells (macroH2A1 KO HFF-T)^22^ with HCMV (*Towne*) and measured infectious viral progeny. We found that HCMV grown in macroH2A1 KO cells produced approximately 30-fold fewer infectious progeny, than HCMV grown in control cells (**Figure 1A, Sup Figure 1A**).

**Figure 1.**
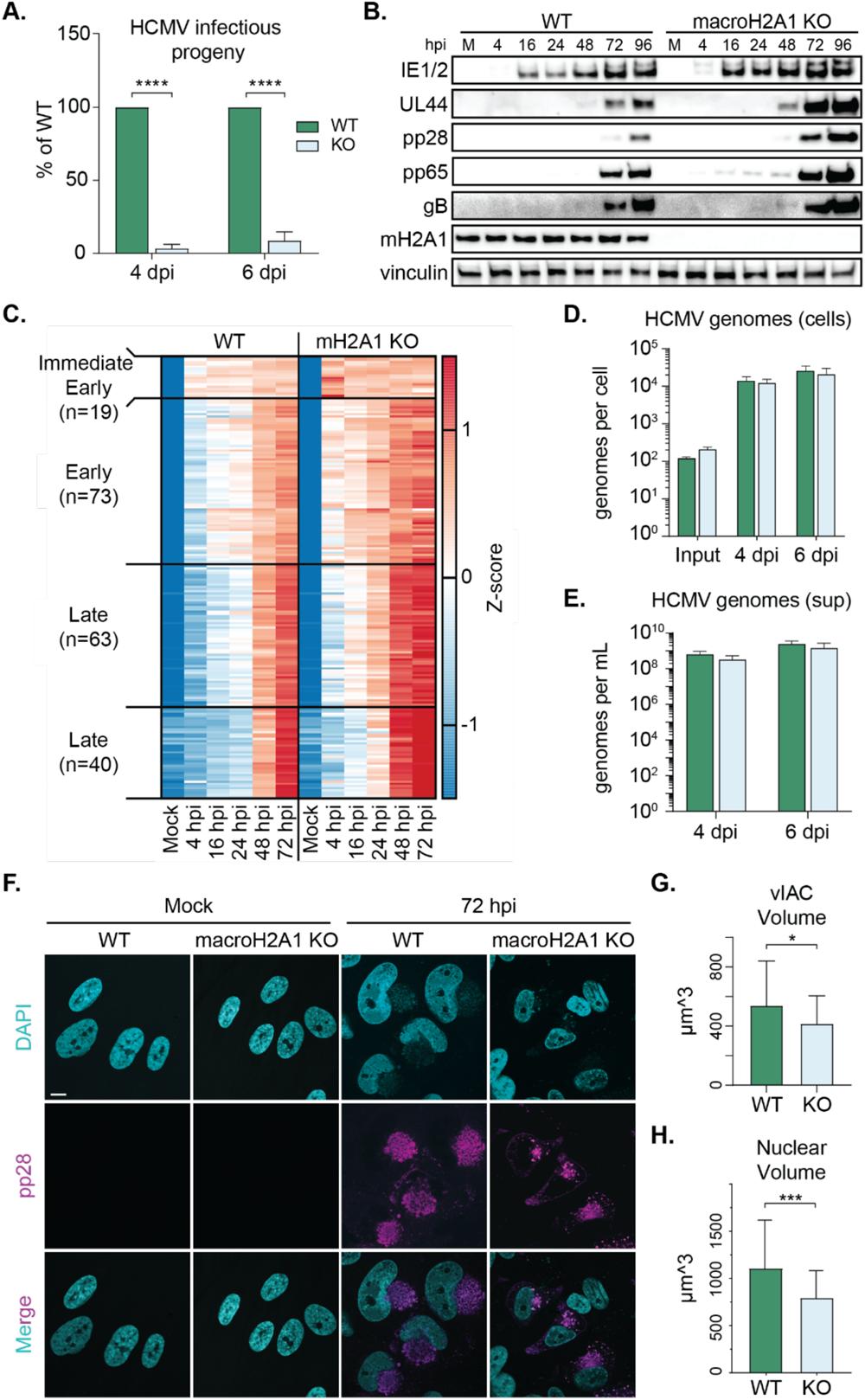
HCMV requires macroH2A1 for efficient production of infectious progeny, but not protein, RNA, or genome accumulation. A) Infectious progeny produced from HCMV infected WT and macroH2A1 KO HFF-T cells quantified by plaque assay at 4 or 6 days post infection (dpi) as indicated. Viral yield is indicated as the percent yield compared to wild type, with errors bars representing SEM. P < 0.0001 at both time points by unpaired T-test. N=3 biological replicates. B) Representative western blots of viral proteins in cells as in (A) during HCMV infection at 4, 16, 24, 48, 72, and 96 hours post infection (hpi) compared to mock (M). These time points correspond to immediate early gene expression (4 hpi), early gene expression (16 hpi), genome replication (24 hpi), and late gene expression (48 and 72 hpi). Vinculin is shown as loading control. C) Heat map of viral genes measured by RNA sequencing at 4, 16, 24, 48, and 72 hpi compared to mock (M). N=3 biological replicates. D) Droplet digital PCR (ddPCR) quantification of HCMV genomes extracted from infected WT and macroH2A1 KO cells at 4 hours (input), 4, and 6 dpi. Bar graphs show the mean with error bars indicating SEM. No significance at any time point by paired T-test. E) ddPCR quantification of HCMV genomes released from cells as in (D) and isolated from supernatants (sups) at 4 and 6 dpi after nuclease treatment, indicating encapsidated genomes. Error bars represent the SEM of three biological replicates. No significance at any time point by paired T-test. F) Representative immunofluorescence images of WT and macroH2A1 KO cells during HCMV infection at mock and 72 hpi. DAPI is shown in cyan, and viral protein pp28 is shown in magenta. Scale bar represents 10 µm. G) Quantification of the volume of viral induced assembly compartments (vIACs) measured by pp28 fluorescence. Bar graphs show mean with error bars indicating SEM. P < 0.05 by unpaired T-test. N>40 vIACs. H) Quantification of nuclear volume of WT and macroH2A1 knockout HCMV-infected cells at 72 hpi. Bar graphs show mean with error bars indicating SEM. p < 0.001 by unpaired T-test. N>40 cells.

We next asked whether macroH2A1 loss affected viral protein and RNA accumulation. We measured viral protein levels by western blot of representative immediate early (IE1/2), early (UL44), late tegument (pp28 and pp65), and late envelope (gB) proteins. We found that all measured viral proteins were robustly expressed at earlier time points, and to a stronger degree in macroH2A1 KO cells (**Figure 1B**).

To determine whether other viral genes not measured by western blot might explain the decrease in infectious progeny produced, we next measured the viral transcriptome by RNA sequencing. We performed bulk RNA sequencing of HCMV-infected WT and macroH2A1 KO cells at 4, 16, 24, 48, and 72 hours post infection (hpi). We found that in macroH2A1 KO cells, early gene expression was initiated by 4 hpi and that immediate early viral genes were expressed at a higher level compared to their levels in WT cells. By 16 hpi, many late viral transcripts were already expressed in macroH2A1 KO cells, while late transcripts were not expressed in WT control cells until 48 hpi (**Figure 1C, Sup Figure 1B-C**). Thus, HCMV transcripts and proteins were expressed earlier and to a higher level in macroH2A1 KO cells compared to WT cells.

As increased protein and RNA expression in macroH2A1 KO cells did not explain the strong reduction in infectious progeny, we next investigated genome replication. We found that within cells, there was no significant change in viral genomes between WT and macroH2A1 KO cells measured by droplet digital PCR (ddPCR) (**Figure 1D**). Similarly, we found no significant change in nuclease-resistant genomes released into the supernatant (**Figure 1E**), indicating that the reduction in infectious progeny is not due to replication or egress defects but rather that viral progeny grown in macroH2A1 KO cells are defective.

### MacroH2A1 is required for nuclear rearrangement and vIAC formation

Proper maturation of HCMV progeny occurs in the vIAC. Therefore, we hypothesized that the loss of macroH2A1 results in a viral maturation defect due to a defective vIAC. We used immunofluorescence to visualize infected cells. Strikingly, we found that the vIACs were significantly smaller in macroH2A1 KO cells (**Figure 1F-G, Sup Figure 1D**), the nuclei did not expand as expected, and the nuclei did not form the characteristic kidney-bean shape (**Figure 1F and H, Sup Figure 1E**). Furthermore, we observed that infected macroH2A1 KO cells also retained the centrosomes and tubulin boundary between cells, suggesting that HCMV-induced syncytia are malformed in infected macroH2A1 KO cells compared to infected WT cells (**Sup Figure 1F**). Our findings demonstrate that macroH2A1 plays a key role in HCMV-induced cellular remodeling and formation of vIACs.

### HCMV rearranges host cytosolic structures in a macroH2A1-dependent manner

To determine if the structural rearrangements that are required for vIAC formation occur in the absence of macroH2A1, we examined cell organization using transmission electron microscopy (TEM). We found that while the overall cellular structure was comparable in mock-treated WT and macroH2A1 KO cells (**Figure 2A**), at 4 days post infection (dpi) HCMV infected cells exhibited dramatically different cytoplasmic structures. We observed that the measurable length of ER regions was significantly shorter in WT cells compared to macroH2A1 KO cells (**Figure 2B-D, Sup Figure 2A-B**). Our findings suggest that without macroH2A1, HCMV is unable to disrupt the host ER.

**Figure 2.**
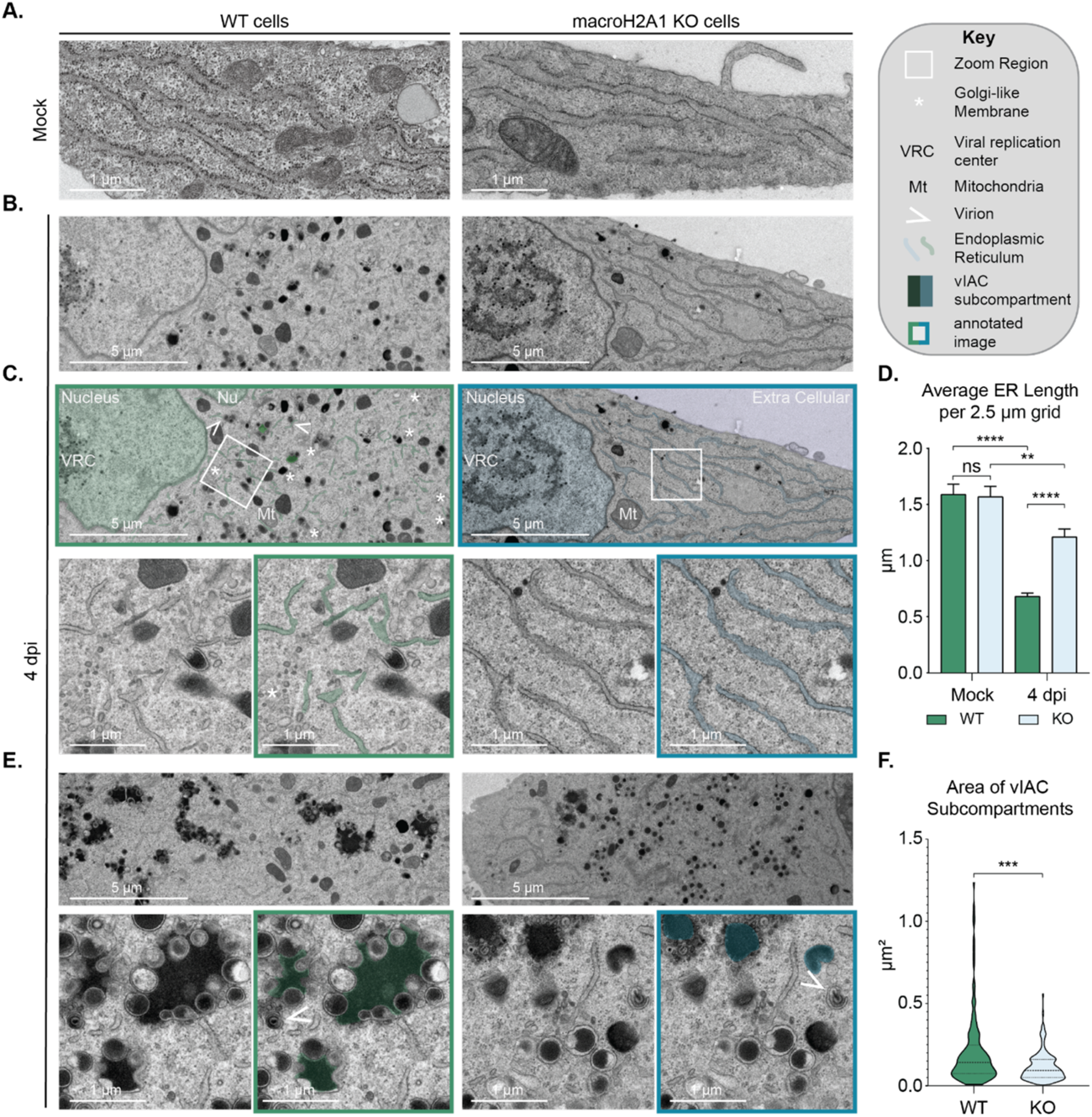
HCMV cellular remodeling and vIAC formation is dependent on macroH2A1. A) Representative transmission electron microscopy images of mock-treated WT and macroH2A1 KO HFF-T cells showing the uninfected state of the endoplasmic reticulum (ER). B) Representative transmission electron microscopy images of WT and macroH2A1 KO cells at 4 days post infection with HCMV. C) Annotation of images from (B). *Right:* Key for annotation. *Below:* Zooms of boxed regions of interest with annotated images of zoomed panels to the right. D) Average endoplasmic reticulum trace per 2.5 µm by 2.5 µm grid of WT and macroH2A1 KO cells in mock and 4 dpi. Bar graph shows mean length of ER per field of view with error bars indicating SEM. ** denotes p < 0.01, **** denotes p < 0.0001 by one-way ANOVA with subsequent Dunnett’s tests of pairs of interest. N=40 grids for mock cells and 100 for infected cells. E) Representative transmission electron microscopy images viral-induced assembly compartments (vIAC) at 4dpi in WT and macroH2A1 KO cells as indicated. *Below:* Zooms from image with annotated versions to the right as in (C). F) Quantification of vIAC subcompartment area. Violin plot depicts median, and upper, and lower quartiles as dotted lines. P < 0.001 by paired T-test. N>100 subcompartments. Scale bars as indicated.

Moreover, by TEM we observed further differences in vIAC formation. In WT cells the vIAC was made up of large subcompartments, consistent with previous findings^24^. Each subcompartment was surrounded by a heterogeneous population of dense bodies, which are non-infectious HCMV particles comprised of enveloped viral proteins^25,26^. In contrast, the HCMV-infected macroH2A1 KO cells rarely formed these distinct large subcompartments (**Figure 2E-F, Sup Figure 2C-D**). In the macroH2A1 KO cells, the largest observable subcompartments, whose area was significantly smaller than those formed in WT cells, were rarely surrounded by dense bodies (**Figure 2E**). The dense bodies in the infected macroH2A1 KO cells were largely homogenous individual structures distributed throughout the cytosol, compared to the heterogenous and conglomerated dense bodies in infected WT cells. Furthermore, virus particles observed in macroH2A1 KO cells frequently appeared malformed, consistent with the finding that macroH2A1 KO cells produce defective progeny (**Figure 2E, arrowhead**). Taken together, these results support our hypothesis that macroH2A1 plays a major role in HCMV-induced cellular remodeling and viral maturation.

### Loss of macroH2A1 prevents activation of neuronal genes during HCMV infection

To profile changes in host gene expression that might lead to the observed phenotypic effects on HCMV infection upon loss of macroH2A1, we analyzed the host transcriptomes of WT and macroH2A1 KO cells during the course of HCMV infection. Principal component analysis of our RNA-seq data showed all replicates clustering close to each other, indicating reproducibility. PC1 captured the time course of infection whereas PC2 captured the genotype (**Sup Figure 3A)**. We first identified a superset of genes that significantly changed in either one of the time points and/or one of the genotypes and then performed k-means (k=4) clustering of Z-scores of gene expression across time and genotype. This analysis captured both time-dependent and genotype-dependent changes in gene expression during HCMV infection (**Figure 3A**). Cluster 1 contained genes that had mixed to low expression in mock-treated cells but steadily increased in expression over the course of infection, peaking at 72 hpi in WT but remained low in the macroH21 KO cells. Genes in this cluster were highly enriched for macroH2A1 and heterochromatin marker H3K27me3 in uninfected cells in our previously published CUT&Tag data set^22^ (**Sup Figure 3B-E**). Clusters 2-4 captured time-dependent changes in expression that were mostly similar between WT and macroH2A1 (**Sup Figure 3F-H**). Cluster 2 contained genes that were activated at 48-72 hpi and expectedly includes genes that would assist in viral trafficking such as those associated with cellular reorganization and protein trafficking within the cell (**Sup Figure 3J**). Cluster 3 contained genes that were activated at 16-24 hpi before returning to low expression. Also as expected, this cluster contains genes associated with immune response and transcription (**Sup.** Figure 3K). Cluster 4 contained genes that were repressed throughout infection and contains genes associated with DNA repair, metabolic processes, and apoptosis (**Sup Figure 3L**). In striking contrast to clusters 2-4, cluster 1 genes were highly enriched for neuronal genes belonging to several Gene Ontology categories that highlighted neuronal function (**Figure 3B, Sup Figure 3I**). Notably, cluster 1 genes strikingly had significantly lower expression in macroH2A1 KO cells, with large fold changes, especially at 72 hpi (**Figure 3C**). Cluster 1 genes, which are the most affected by the loss of macroH2A1, suggest that HCMV induces a neuronal-like transcriptional profile that increases over the course of infection.

**Figure 3.**
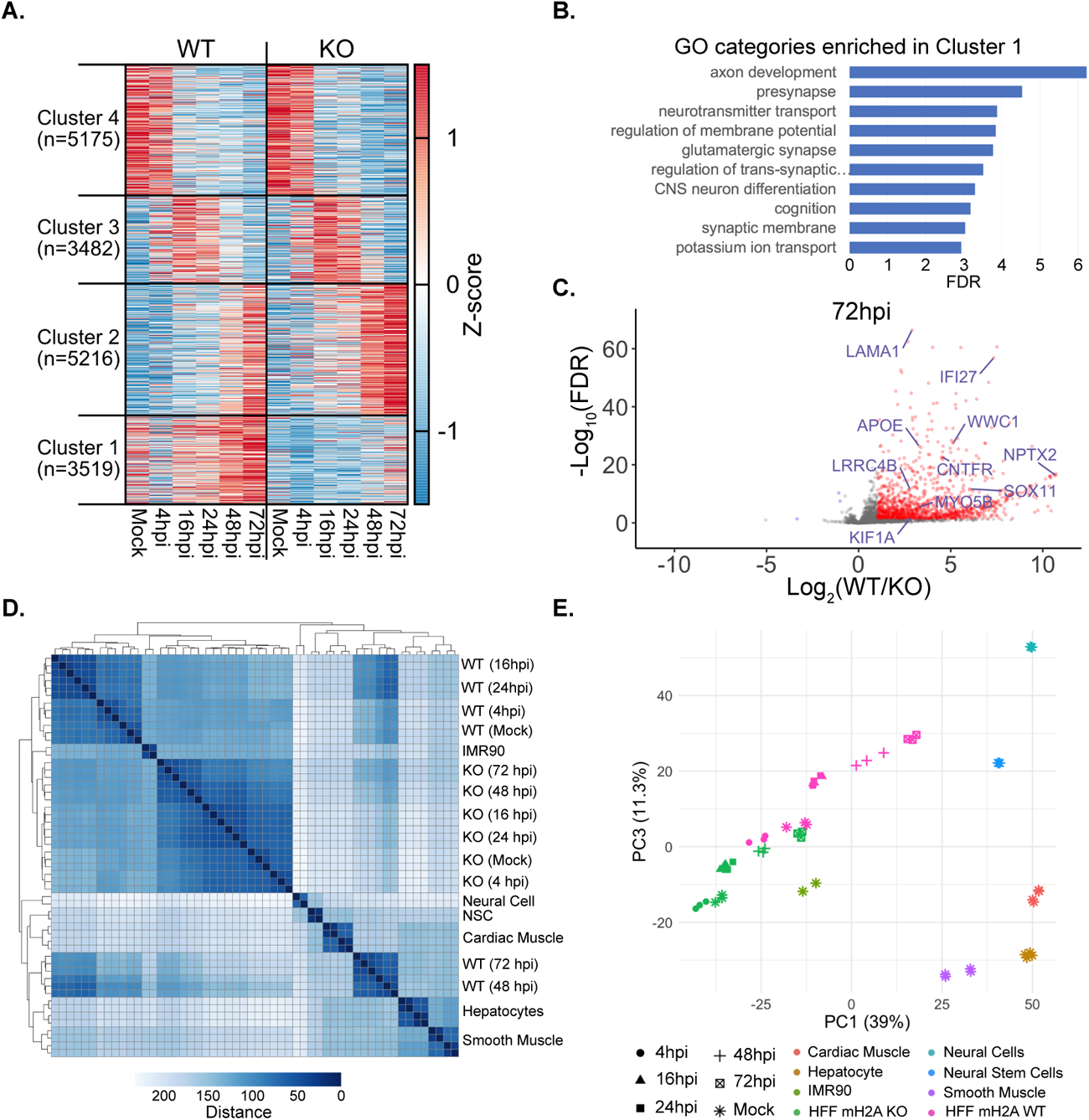
Host gene expression is altered upon loss of macroH2A1 during HCMV infection. A) K-means clustering (k=4) of gene expression changes over 72 hours of infection shown as a heatmap. Z-scores were calculated for each gene from its normalized count across the time course of CMV infection for WT and macroH2A1 KO cells. B) The -log_10_(FDR) value for enrichment of neuronal GO categories in Cluster 1. C) Volcano plot where the Log_2_(Fold Change) for WT vs. macroH2A.1 KO is plotted against -log_10_(FDR) for genes in Cluster 1. Genes with Log_2_(Fold Change) >1 and FDR ≤ 0.05 are marked in red. Neuronal genes selected for further characterization are labeled in blue. D) Matrix of Euclidean distance between normalized expression profiles of CMV infection time course for WT and macroH2A1 KO, and other cell types. Gene expression datasets for other cell types were obtained from ENCODE. E) PCA plot showing PC1 and PC3 for the same expression profiles plotted in (D).

To test the hypothesis that HCMV induces a neuronal-like phenotype that is blunted by the loss of macroH2A1, we compared our gene expression profiles to those of different cell types from ENCODE^27,28^. From ENCODE, we included expression profiles of IMR90 fibroblast cells, similar to our HFF cells, and cells differentiated from induced pluripotent stem cells to form cells in the neuronal, muscle, and liver lineages. We first generated a distance matrix and observed that IMR90 clustered with all macroH2A1 KO time points and WT mock and early time points. However, WT cells at 48 and 72 hpi cluster with the other lineages we included in the analysis (**Figure 3D**). This suggests that later time points of HCMV infection cause a transition of HFF cells to a non-fibroblast identity, and this transition is suppressed in the absence of macroH2A1. To explore this further, we performed a principal component analysis of the expression matrix comprising genes from Cluster 1. PC1 captures the differences between fibroblast and non-fibroblast lineages, whereas PC3 captures the differences between neuronal and non-neuronal lineages (**Figure 3E**). We found PC1 and PC3 to capture the infection time course. WT mock-infected cells had similar values to IMR90 in PC1 and PC3. We observed an increase in PC1 and PC3 loadings with increasing time of infection in the direction toward neural cells. A similar trend is observed in macroH2A1 KO cells, but the starting time points (mock and 4 hpi) have much lower PC1 and PC3 loadings such that by the final time point, PC1 and PC3 values for macroH2A1 KO cells are similar to that of mock-infected WT cells. Thus, an increase in both PC1 and PC3 observed over the course of infection captures the loss of fibroblast identity and a gain of neuronal identity in WT cells. Interestingly, in macroH2A1 KO cells, the starting point is much lower in PC1 and PC3, suggesting an inability of macroH2A1 KO cells to transition from fibroblast to neuronal identity over the course of infection. In summary, our gene expression analysis highlights the profound effect of macroH2A1 on transcriptional upregulation of many host genes during HCMV infection and that this upregulation transitions infected cells away from a fibroblast expression profile and towards a neuronal expression profile.

As macroH2A1 frequently colocalizes with H3K27me3 and cluster 1 was also enriched for H3K27me3 (**Sup Figure 3D-E**), we used the small molecule tazemetostat (an EZH2 inhibitor)^29^, to deplete H3K27me3 prior to HCMV infection. We found that although H3K27me3 depletion caused a significant reduction in titer, the reduction was modest compared to that induced by loss of macroH2A1 (**Sup Figure 4A**). Furthermore, H3K27me3 depletion did not impact nuclear rearrangement or vIAC formation (**Sup Figure 4B-D**). We also investigated by western blot the induction of cluster 1 gene KIF1A, a kinesin motor for axonal transport in neurons^30^, and found that its induction was not diminished significantly by the depletion of H3K27me3 (**Sup Figure 4E**). Our results suggest that while H3K27me3 is likely also important for HCMV infection, its role on cluster 1 genes appears less significant for HCMV infection than that of macroH2A1.

### Knockdown of several neuronal genes reduced HCMV spread and vIAC formation

We next sought to determine if the transition from a fibroblast to neuronal expression profile is essential for HCMV infectivity. Following the observation that many of the genes differentially expressed in HCMV infection between WT and macroH2A1 KO cells were associated with axon formation and neurotransmitter trafficking, we hypothesized that HCMV hijacks these pathways for progeny maturation and spread. To test this hypothesis, we designed a targeted siRNA screen selecting 12 genes with low FDRs and large fold changes in expression between WT and macroH2A1 KO cells induced by HCMV at 72 hpi (**Sup Figure 5A-C**).

Following infection of WT cells with HCMV tagged with GFP, we transfected siRNAs to dampen the induction of these target genes later in infection (**Figure 4A**). We confirmed that all siRNA targeted transcripts were reduced by at least 50% compared to their levels at 4 dpi in the condition treated with a non-targeting control (NC) siRNA (**Figure 4B Sup Figure 5D**). Upon initial infection, representative HCMV RNA and protein levels did not differ among any conditions (**Sup Figure 5E-F**). This indicates viral transcription and translation were not impacted by the siRNA treatment. We used supernatants harvested from the siRNA-treated HCMV-infected cells to set up GFP foci assays and measure plaque size. We found that several siRNA knockdowns caused a reduction in GFP foci to 40-70% of control levels though these groups did not reach significance (**Figure 4C**). Knockdown of WWC1, however, did significantly reduce GFP foci. Interestingly, we also noted that several of the knockdown conditions produced viral progeny that generated smaller plaque sizes compared to the control. To investigate this observation, we imaged crystal violet plaques and quantified the area of these plaques (**Sup Figure 6A**). We found that five of our target genes, IFI27, KIF1A, LAMA1, NPTX2, and WWC1, had significantly smaller plaques compared to control (**Figure 4D-E**).

**Figure 4.**
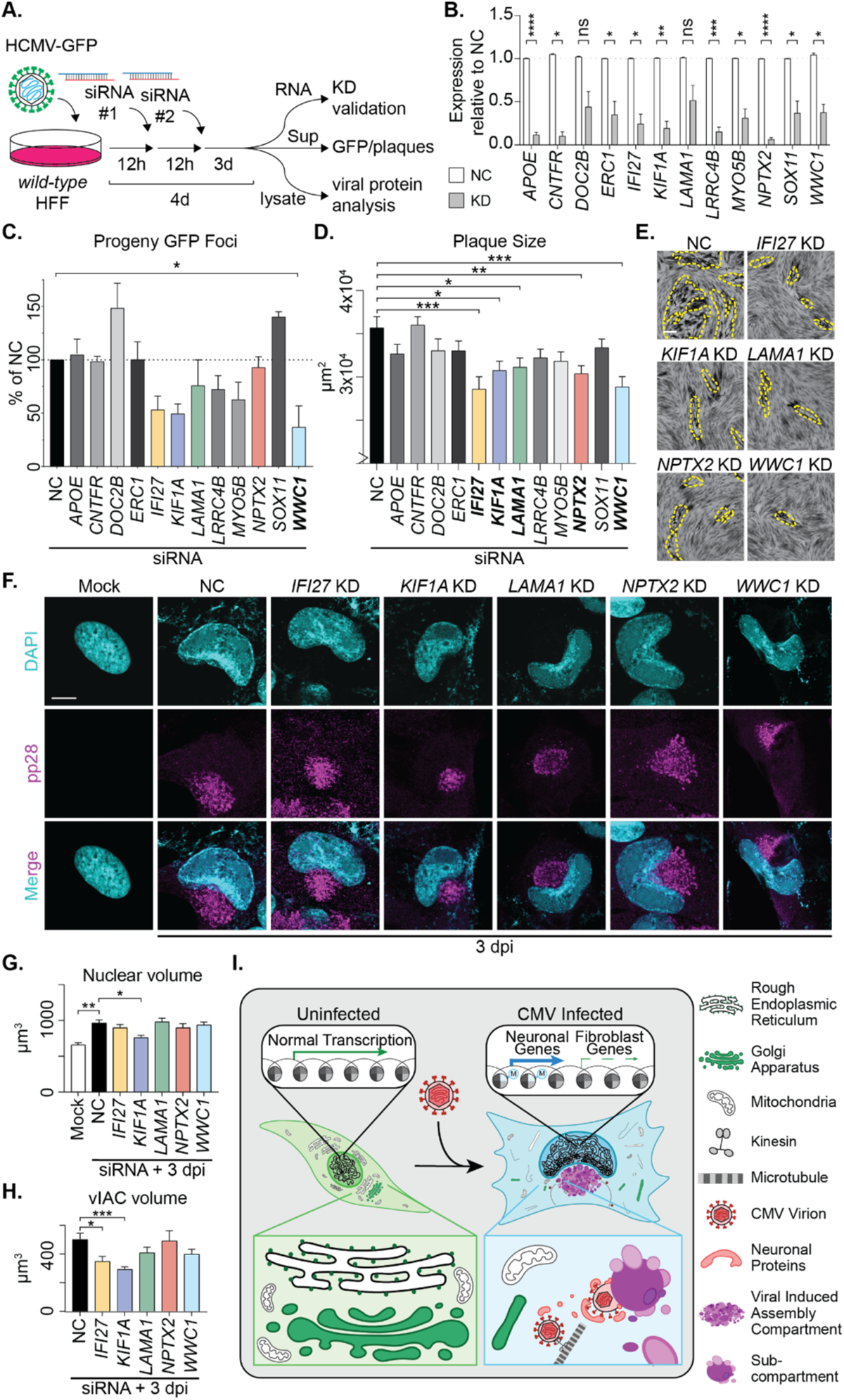
Successful HCMV maturation requires induction of dormant neuronal proteins. A) Schematic of targeted siRNA screen methodology. B) RT-qPCR quantification of RNA levels of target genes during HCMV infection. These genes include *APOE*, a lipoprotein associated with Alzheimer’s disease and synaptic vesicle release^41,42^; *CNTFR*, a ciliary neurotrophic factor receptor that supports motor neuron axons^43^; *DOC2B* a calcium sensor that promotes synaptic vesicle release^44^; *ERC1*, a cellular scaffolding protein^45^; *IFI27*, an interferon-induced gene expressed in the cerebellum in response to viral CNS infection^46^; *KIF1A*, a neuronal kinesin^30^; *LAMA1*, a laminin essential for neurite growth^47,48^; *LRRC4B*, a transmembrane protein that regulates synapse formation^49^; *MYO5B*, a myosin associated with polarity and axon development^50^; *NPTX2* (formerly *NARP*), a small molecule released in excitatory synapses^51^; *SOX11*, a transcription factor associated with neuron development^52^; and *WWC1*, a synaptic scaffolding protein^53^. Knockdown of each gene at 4dpi is normalized to its expression in cells treated with the non-targeting control (NC) at 4 dpi. Bar graphs show mean with error bars indicating SEM. N=3 biological replicates. C) Quantification of GFP foci in cells infected with supernatant harvested from cells treated with indicated siRNA as depicted in (A). Bar graphs show mean with error bars indicating SEM. * indicates P < 0.05 by one way ANOVA with follow up Dunnett’s test. N=3 biological replicates. D) Quantification of plaque area produced from supernatant harvested from cells treated with siRNA indicated. Those that reach statistical significance are bolded. Bar graphs show mean with error bars indicating SEM. * denotes P < 0.05, ** denotes p < 0.01, *** denotes p < 0.001 by one way ANOVA with follow up Dunnett’s test. N > 300 plaques. E) Representative images of plaque sizes for those with significant differences as indicated. Yellow dashed line frames plaque example. Scale bar indicates 150 µm. F) Representative immunofluorescence images of HCMV-infected cells treated with indicated siRNA knockdown. DAPI is shown in cyan and pp28 is shown in magenta. Scale bar represents 10 µm. G) Quantification of nuclear volume in siRNA-treated cells infected with HCMV. Bar graph shows mean with error bars indicating SEM. * denotes P < 0.05, ** denotes p < 0.01 by one way ANOVA with follow up Dunnett’s test. N > 60 cells. H) Quantification of viral induced assembly compartment (vIAC) volume in siRNA-treated cells infected with HCMV. Bar graph shows mean with error bars indicating SEM. * denotes P < 0.05, *** denotes p < 0.001 by one way ANOVA with follow up Dunnett’s test. N > 60 vIACs. I) Model schematic. HCMV-infected cells upregulate numerous neuronal genes and these genes are required by the virus for proper cellular remodeling, formation of the viral assembly compartment, and viral maturation to promote viral spread.

Next, we investigated whether depletion of these five targets impacted HCMV-induced cellular remodeling and vIAC formation. We found that IFI27 and KIF1A KD cells had smaller nuclei and significantly smaller vIACs compared to control. Additionally, LAMA1, NPTX2, and WWC1 KD resulted in vIACs that were malformed with either a hollow or nonspherical appearance (**Figure 4F-H and Sup Figure 6B-D**). Taken together, our results from this screen demonstrate that cells unable to induce these genes to a high level are unable to produce viral progeny that can spread efficiently, underscoring the importance of these genes in HCMV spread.

## Discussion

In this study, we demonstrated that human cytomegalovirus induces expression of numerous neuronal genes involved in synapse formation and neurotransmitter vesicle trafficking in a macroH2A1-dependent manner. We further showed that macroH2A1 and several of these induced neuronal genes are crucial to HCMV maturation and spread (**Figure 4I**). Our findings indicate that viral reprogramming of the host cell is dependent on host chromatin-controlled changes and uncovers previously unappreciated pathway critical for HCMV infection.

The formation of a vIAC and kidney-bean shaped nucleus was thought to be an exclusive feature of HCMV infection, however, these changes have also been observed in HSV-1 infection of neuron-like cells^31^. In fact, one of our identified neuronal genes important for HCMV maturation, *KIF1A*, a kinesin motor protein involved in axonal transport, was shown to be important for the spread of HSV-1 and pseudorabies virus (PRV)^32^. These observations raise an interesting question about whether HCMV hijacking of neurotransmitter trafficking pathways is a retained evolutionary feature of many viruses or a novel pathway exploited specifically by HCMV. MacroH2A1 is also highly conserved and not rapidly evolving^33^, suggesting that it is more likely to be HCMV that evolved to hijack this histone. Future work into the evolution of HCMV will uncover how these mechanisms of chromatin manipulation have developed.

Murine CMV was recently reported to control large scale transcriptional profiles and alter the identity of infected macrophages to escape innate immune response and increase spread^34^. Our work builds on this finding in HCMV, suggesting that betaherpesviruses may drive cellular reprogramming for infection spread in various cell-type specific ways. Importantly, we find primarily structural genes associated with terminal differentiation to be upregulated during HCMV infection, as opposed to developmental genes. Taken together with the recent findings on murine CMV, our work suggests that HCMV cellular reprogramming is not limited to a particular set of genes, but rather is controlled through chromatin mechanisms. Furthermore, our discovery that HCMV upregulates genes associated with synapse formation and neurotransmitter trafficking for their efficient spread provides a functional context for the previous findings wherein HCMV virions resemble synaptic-like vesicles in their lipid content^35^.

It is important to note that these neurotransmitter pathways are dormant in uninfected fibroblast cells. Thus, there are no normal neuronal secretory functions that the virus must compete with, nor are there specific neuronal immune defenses to subvert. Furthermore, neuronal trafficking is one of the fastest and most efficient mechanisms by which to exit the cell^36^. Therefore, HCMV has pinpointed and activated an entirely dormant pathway to both avoid competition and successfully egress from the infected cell. Interestingly, the idea that ancient human retrovirus integration gave rise to the neuronal trafficking protein, Arc^37^, further supports the hypothesis that viral and neurotransmitter trafficking rely on similar mechanisms.

Our study has revealed how HCMV remodels the host transcriptome in a macroH2A1-dependent manner to promote viral maturation and spread. Future work to identify the viral mechanism that drives this macroH2A1-dependent cellular reprogramming is expected to shed light on many new questions arising from this study. Extensive previous work demonstrated that the HCMV protein IE1/2 can directly bind the acid patch of the nucleosome to affect chromatin^38,39^, however, it is unknown whether IE1/2 interacts with macroH2A1-containing nucleosomes. New screening methods may also uncover unknown strategies for chromatin manipulation through other factors to promote viral maturation^40^, though targeted approaches for chromatin may be required. Given our findings that HCMV induces expression of neuronal genes through chromatin manipulation, it is likely that many additional mechanisms of chromatin hijacking by viruses have yet to be uncovered.

## Author Contributions

LEKM-Conceptualization, Methodology, Formal analysis, Investigation, Data Curation, Writing-Original Draft, Writing-Review and Editing, Visualization, Supervision. JRS-Conceptualization, Methodology, Formal analysis, Investigation, Data Curation, Writing-Review and Editing, Visualization. HCL-Conceptualization, Investigation, Writing-Review and Editing. LSW-Investigation, Data Curation, Writing-Review and Editing. DHN-Investigation, Data Curation, Writing-Review and Editing, Visualization. EAA-Investigation, Writing-Review and Editing.

MRB-Investigation, Writing-Review and Editing. APG-Conceptualization, Methodology, Writing-Review and Editing, Resources, Supervision. SR-Conceptualization, Methodology, Formal analysis, Software, Data Curation, Writing-Original Draft, Writing-Review and Editing, Visualization, Supervision. DCA-Conceptualization, Methodology, Formal analysis, Investigation, Resources, Data Curation, Writing-Original Draft, Writing-Review and Editing, Visualization, Supervision, Funding acquisition.

## Acknowledgements

We thank members of the Avgousti lab, M. Lagunoff, S. Parkhurst, S. Tapscott, L. Goo and M. Weitzman for their insightful comments. We thank the Electron Microscopy (EMSR), and Genomics and Bioinformatics Shared Resource Facilities at the Fred Hutchinson Cancer Center for help with sequencing and data analysis. We thank H. West-Foyle, L. Schroeder, and the Cellular Imaging Shared Resource (CISR) for their help with image analysis.

This research was supported by the Electron Microscopy Shared Resource, RRID:SCR_022611, The EMSR, CISR, and Genomics core are supported in part by the Fred Hutch/University of Washington Cancer Consortium (P30 CA015704). This study was also supported by start-up funds from the RNA Bioscience Initiative at the University of Colorado School of Medicine (S. Ramachandran), the Fred Hutch (D.C. Avgousti), National Institutes of Health funding to E.A. Arnold (AI083203), A.P. Geballe (AI145945), S. Ramachandran (GM133434), and D.C. Avgousti (GM133441), and the University of Washington Magnuson Scholarship to L.E. Kelnhofer-Millevolte.

## Supplemental Figures

**Figure S1.**
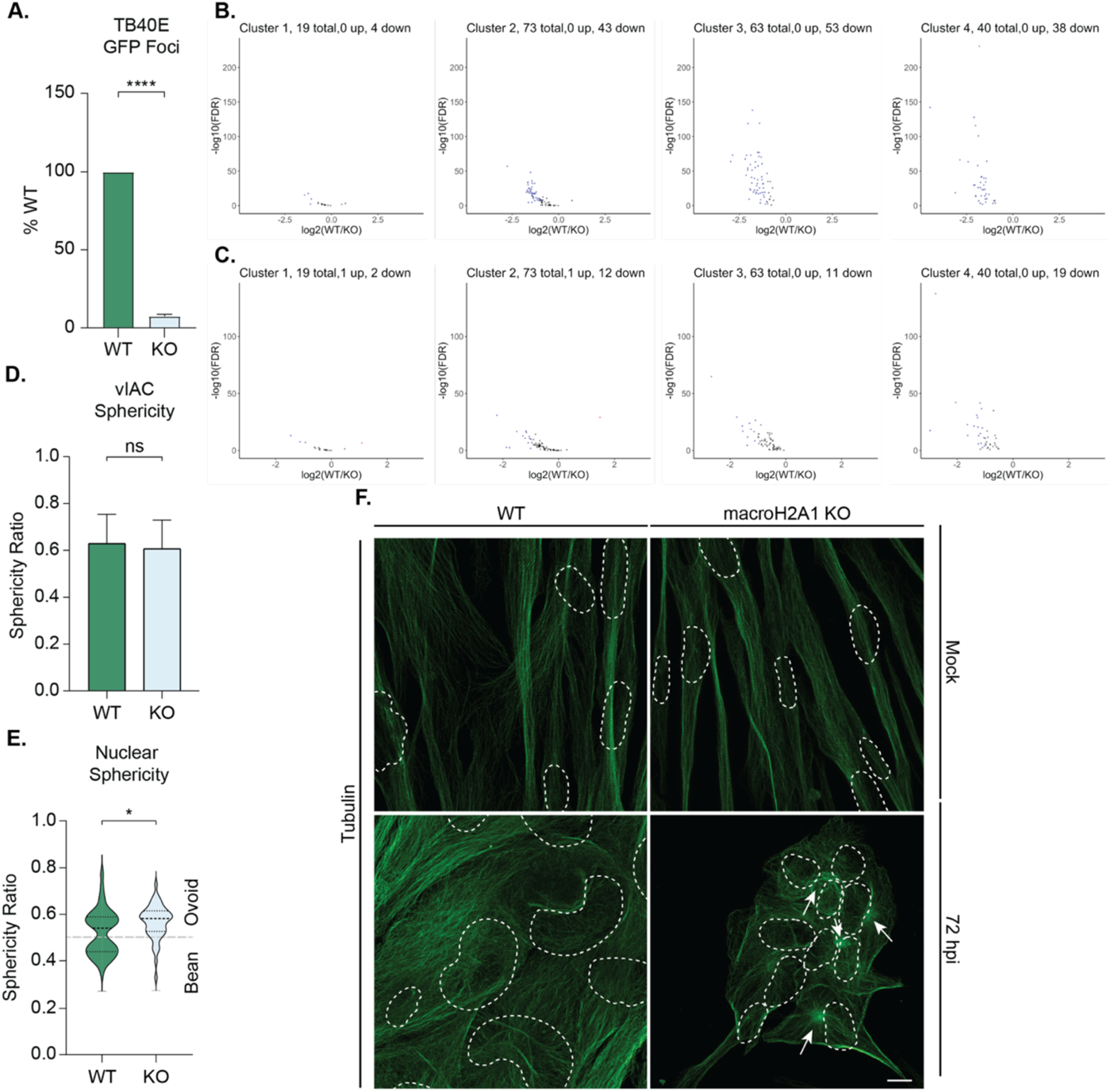
HCMV nuclear and cytoskeletal reorganization requires macroH2A1. A) Infectious progeny produced from ***TB40E-GFP*** HCMV infected WT and macroH2A1 KO HFF-T cells quantified by plaque assay at 4 or 6 days post infection (dpi) as indicated. Viral yield is indicated as the percent yield compared to wild type, with errors bars representing SEM. P < 0.0001 at both time points by unpaired T-test. N=3 biological replicates. B) Volcano plots where the Log_2_(Fold Change) for WT vs. macroH2A1 KO at 48 hpi is plotted against -log_10_(FDR) for genes in clusters as marked. Significantly upregulated genes (Genes with Log2(Fold Change) >1 and FDRσ≤0.05) are marked in red, and significantly downregulated genes (Genes with Log2(Fold Change) <1 and FDRσ≤0.05) are marked in blue. C) Volcano plots as in (B) for gene expression at 72 hpi. D) Sphericity of viral induced assembly compartment as measured by pp28 staining. Bar graphs show mean with error bars indicating SEM. No significance by unpaired T-test. N>40 vIACs. E) Nuclear sphericity of WT and macroH2A1 KO HCMV infected cells at 72 hpi. Violin plot depicts median and quartiles in dashed lines. Division of oval and bean shape as marked. p < 0.05 by unpaired T-test. N > 40 cells. F) Representative immunofluorescence images of WT and macroH2A1 KO cells during CMV infection at mock and 72 hpi with tubulin staining in green. Arrows indicate centrosomes and dashed white lines outline nuclei. Scale bar represents 10 micrometers.

**Figure S2.**
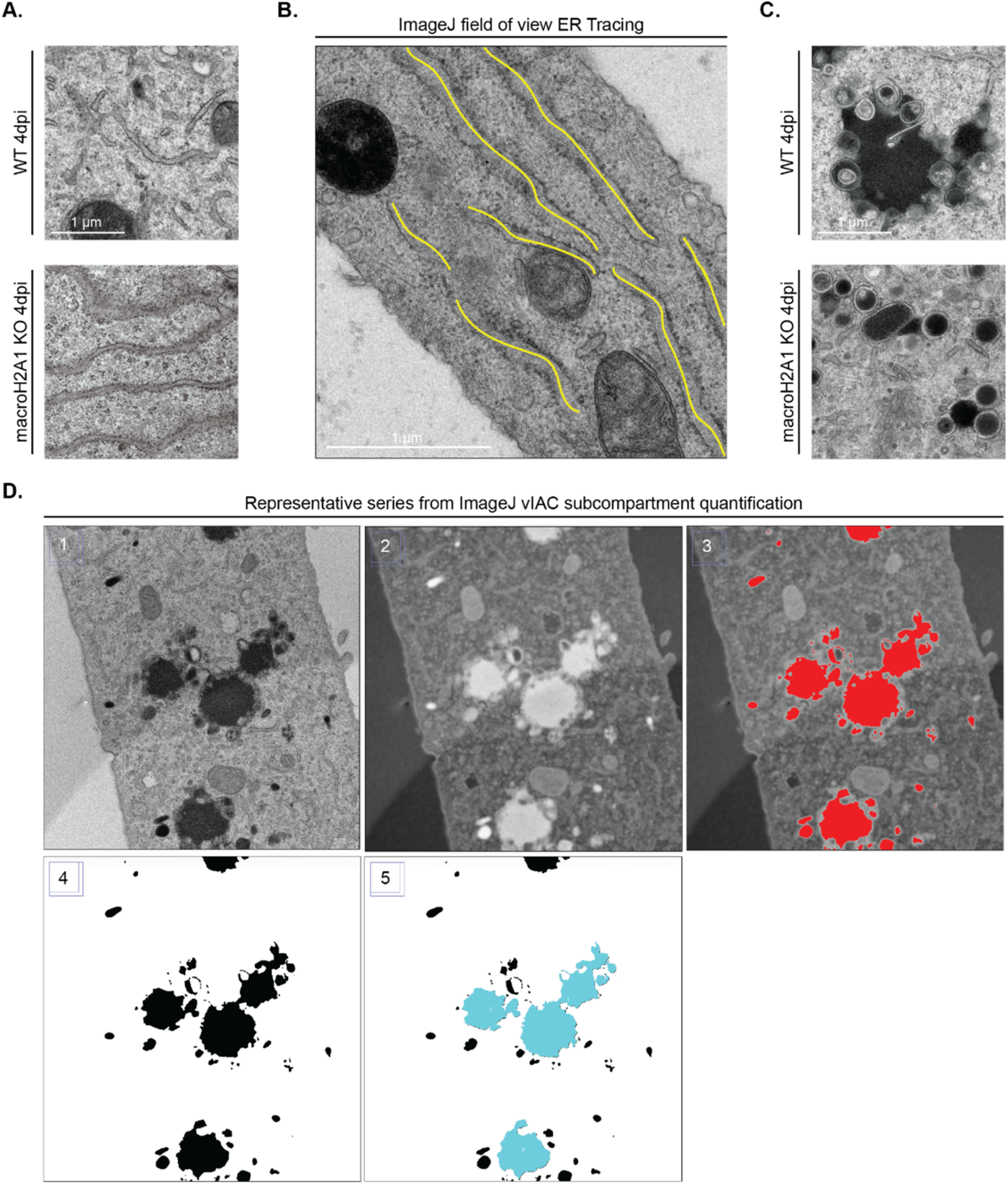
Image analysis pipeline for electron microscopy data. A) Additional transmission electron microscopy image of WT and macroH2A1 KO cells at 4 dpi with HCMV. Scale bar as indicated. B) Representative image highlighting the ER tracing in ImageJ for quantification in Figure 2D. Yellow lines represent traces made with “Freehand line tool”, length of line was measured with ImageJ “Measure” function. C) Additional transmission electron microscopy images of vIACs from WT and macroH2A1 KO cells at 4 dpi. D) Representative images indicating the ImageJ macro for quantification shown in Figure 2F. 1) Original image, 2) invert and gaussian blur, 3) thresholding to define subcompartments, 4) convert to mask and fill holes, 5) size exclusion on dense bodies and virions.

**Figure S3.**
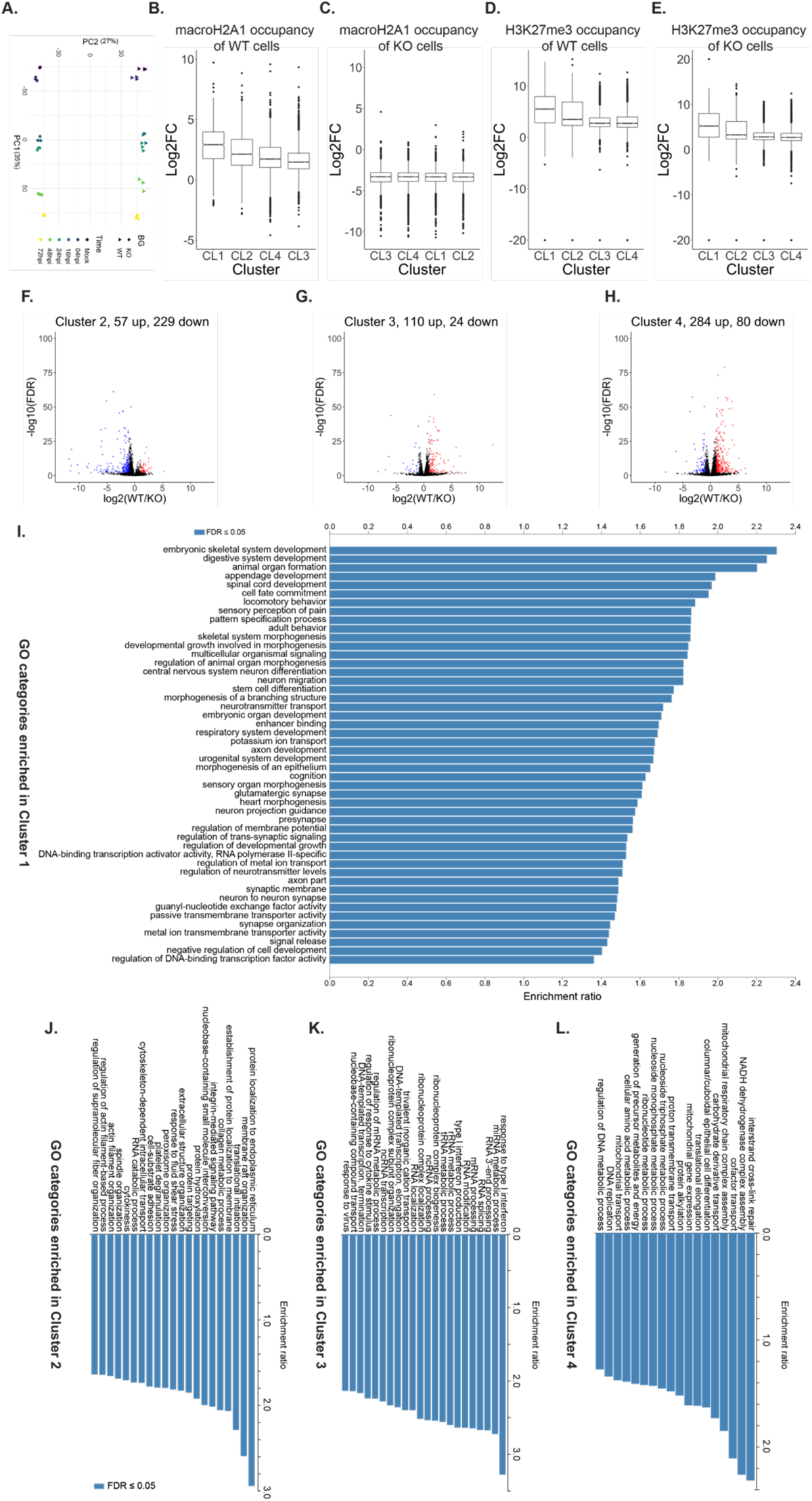
Host gene expression and chromatin states. A) Loadings of first two principal components from principal component analysis of the matrix of quantification of all genes for all 36 samples. PC1 captures the time course of infection, whereas PC2 captures the genotype. B) Log2 fold change of macroH2A CUT&Tag enrichment compared to IgG from uninfected cells at genes (from gene start to gene end) in each cluster plotted as boxplots. C) Same as (B) for macroH2A1 KO cells. D) Same as (B) for H3K27me3 CUT&Tag in WT cells. E) Same as (B) for H3K27me3 for macroH2A1 KO cells. Published macroH2A, H3K27me3, and IgG CUT&Tag data were used for (B-E). F-H) Volcano plots where the Log_2_(Fold Change) for WT vs. macroH2A.1 KO at 72 hpi is plotted against -log_10_(FDR) for genes in clusters 2 (F), 3(G), and 4 (H) shown. Significantly upregulated genes (Genes with Log2(Fold Change) >1 and FDR≤0.05) are marked in red, and significantly downregulated genes (Genes with Log2(Fold Change) <1 and FDR≤0.05) are marked in blue. I-L) Enrichment of GO categories with FDR<0.05 plotted for Cluster 1 (I), 2 (J), 3(K), and 4 (L).

**Figure S4.**
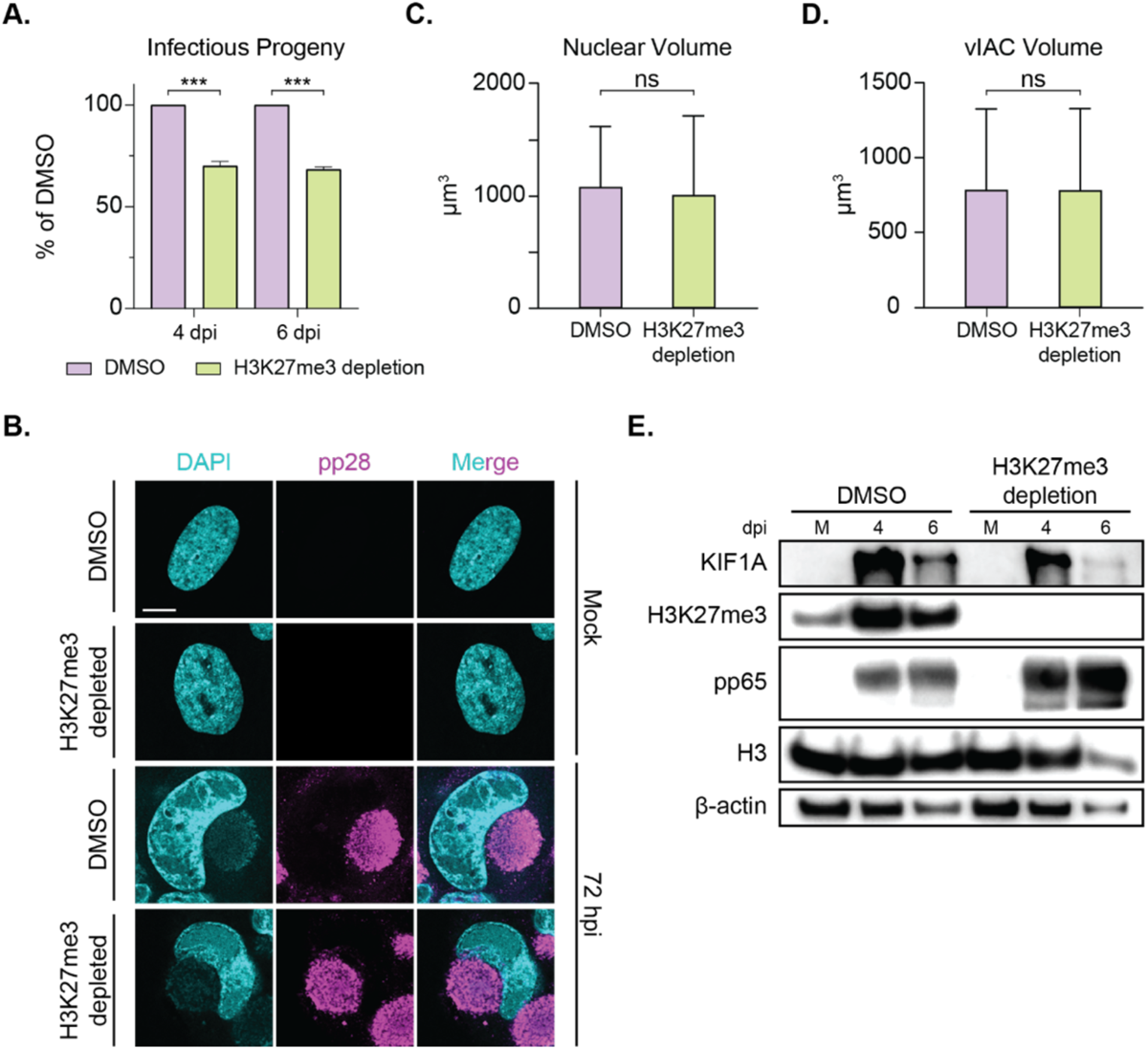
H3K27me3 is not required for cellular remodeling or KIF1A induction by HCMV. A) Infectious progeny produced from HCMV-infected WT and H3K27me3 depletion by tazemetostat cells quantified by plaque assay. Viral yield is indicated as the percent yield compared to wild type, with errors bars representing SEM. P < 0.001 at both time points by unpaired T-test. N=3 biological replicates at 4 dpi N=2 at 6 dpi. B) Representative immunofluorescence images of WT and H3K27me3 depleted cells during HCMV infection at mock and 72 hpi. DAPI is shown in cyan and viral protein pp28 is shown in magenta. Images are merged in bottom row. Scale bar represents 10 µm. C) Nuclear volume of WT and H3K27me3-depleted HCMV-infected cells at 72 hpi. Bar graphs show mean with error bars indicating SEM. No significance by unpaired T-test. N > 40 cells. D) Volume of viral induced assembly compartment as measured by pp28 staining. Bar graphs indicate mean with error bars indicating SEM. No significance by unpaired T-test. N > 40 vIACs. E) Representative western blot of viral proteins in WT and H3K27me3 depleted cells during HCMV infection at 4 or 6dpi compared to mock (M) as indicated. Vinculin is shown as loading control.

**Figure S5.**
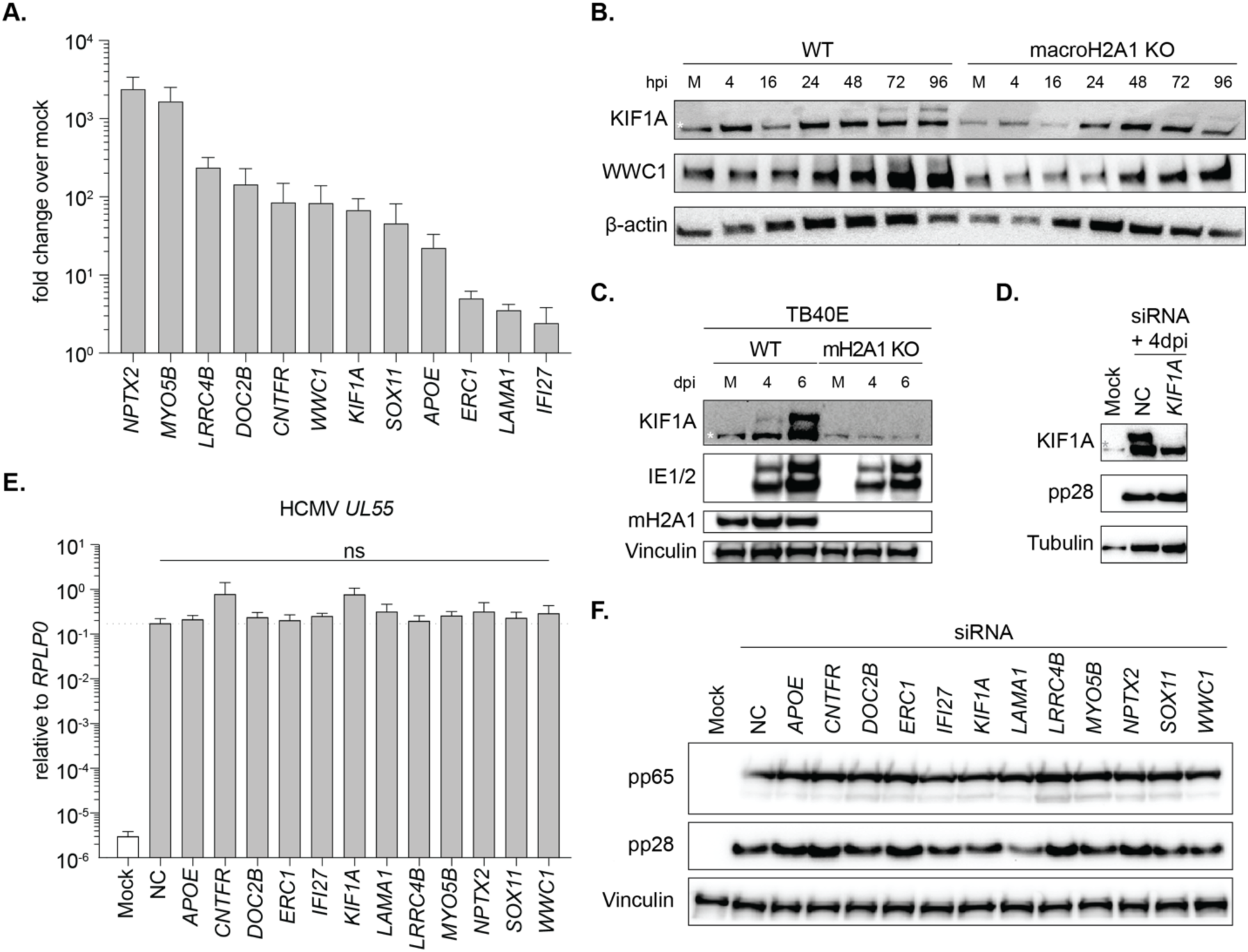
Gene expression analysis of siRNA screen during HCMV infection. A) RT-qPCR of target gene RNA levels during HCMV infection at 4 dpi. Bar graphs indicate mean with error bars indicating SEM. N=3 biological replicates. B) Representative western blot of neuronal proteins in WT and macroH2A1 KO cells during HCMV infection at 4, 16, 24, 48, 72, and 96 hpi compared to mock (M). Asterisk indicates a non-specific band. Actin is shown as loading control. C) Representative western blot of KIF1A in WT and macroH2A1 KO cells during *TB40E-GFP* HCMV infection at 4 and 6 dpi compared to mock (M). Asterisk indicates a non-specific band. Vinculin is shown as loading control. D) Representative western blot of siRNA knockdown in WT cells during HCMV infection at 4 dpi compared to mock (M). Asterisk indicates a non-specific band. Tubulin is shown as loading control and pp28 is shown as infection control. E) RT-qPCR of HCMV UL55 in siRNA treated cells during CMV infection. Expression is normalized to the non-targeting control treated mock-infected cells. Bar graphs show mean with error bars indicating SEM. N=3 biological replicates. No significance by ANOVA. F) Representative western blot of viral proteins pp65 and pp28 in siRNA knockdown in WT cells during HCMV infection at 4 dpi compared to mock (M). Vinculin is shown as loading control.

**Figure S6.**
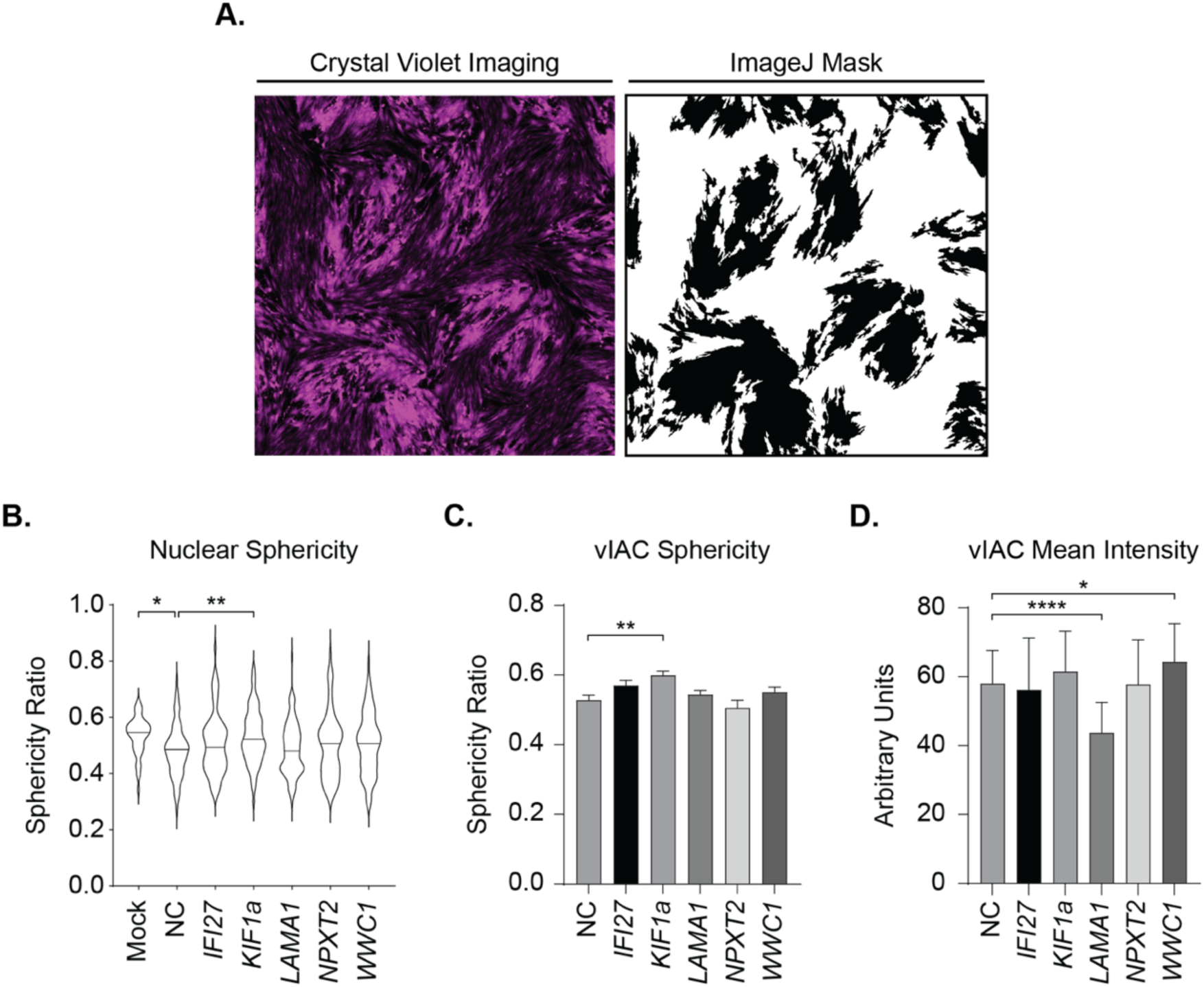
Image analysis of siRNA screen. A) Representative Cy-5 imaging of Crystal Violet stained HCMV plaques in HFF cells and subsequent ImageJ quantification. B) Nuclear sphericity of HCMV-infected WT and siRNA treated cells as indicated at 72 hpi. Violin plot depicts median and quartiles in dashed lines. * indicates p < 0.05 ANOVA with follow up Dunnett’s test. N > 40 cells. C) Sphericity of viral-induced assembly compartment as measured by pp28 staining. Bar graphs show mean with error bars indicating SEM. No significance by unpaired T-test. N > 40 vIACs. D) Average intensity of viral-induced assembly compartments as measured by pp28 staining. Bar graphs show mean with error bars indicating SEM. * indicates p < 0.05 ANOVA with follow up Dunnett’s test. N > 40 vIACs.

## Material and Methods

### Cells and viruses

hTERT-immortalized HFFs, and hTERT-immortalized macroH2A1 knockout HFFs generated as previously described^22^, were cultured using standard methods with 10% FBS and 1% penicillin-streptomycin as previously described^54^. Cells were grown at 37°C with 5% CO_2_ and routinely tested for mycoplasma contamination.

The lab-adapted strain of HCMV *Towne*^55^ was used for all experiments unless otherwise noted. GFP-*Towne*^56^ and *TB40E*-GFP^57^ were used for experiments indicated at an MOI of 1. Monolayers of cells were infected for 1 h at 37°C as previously described^58^. Cells were collected at 4, 16, 24, 48,72, 96 hpi for western blot and RNA-sequencing. The supernatant was collected at 4 and 6 dpi for plaque assays. Virus stock was grown by infecting WT HFF cells at an MOI of 0.0001. The virus was harvested ∼16-20 dpi and titered on HFF cells to determine stock plaque-forming units per ml (PFU/ml). Experimental plaque assays were set up in WT HFF cells. Plaque assays were set up as serial 10-fold dilutions in serum-free DMEM. The virus was left on the cells for 1 h and then aspirated. Cells were washed with 1× PBS (pH 7.46) and 2% methylcellulose overlay in DMEM with 2% FBS, and 0.5% penicillin-streptomycin was added to wells. Plaques were fixed with 0.2% crystal violet at 14 dpi and plaques were counted by hand. All plaque assays were set up with two technical replicates.

In the case of GFP-tagged viruses, foci were read at 7 dpi using the Cy-5 filter on a Typhoon Trio Imager and quantified using FIJI is Just ImageJ version 2.1.0: Java 1.8.0_172 [64-bit].

### Infections with tazemetostat pretreatment

HFFs were treated with DMSO or 10 µM of tazemetostat (HY-13803; MedChem) in DMSO for 3 d prior to infection as previously described^22^. Cells were then infected at an MOI of 1, and after 1 h of incubation with the virus, fresh media with 10 µM tazemetostat was added to previously treated cells. Control samples were treated with equivalent volumes of DMSO. Samples were harvested as above.

### Western blotting

Western blotting was performed as previously described^54^. Briefly, cells were counted, pelleted, resuspended in 1× NuPAGE lithium dodecyl sulfate (LDS) sample buffer (NP007; Thermo Fisher Scientific) + 5% β-mercaptoethanol at 300,000 cells per 200 µl, and boiled for 15 min.

Protein lysates were separated by 13.5% SDS-PAGE gels using 1× NuPAGE MOPS buffer (NP0001; Thermo Fisher Scientific) at 75 V for 30 min, then 110 V for 100 min, and then wet transferred to a nitrocellulose membrane (Bio-Rad) at 100 V for 70 min using Transfer Buffer (25 mM Tris Base, 100 mM glycine, 20% methanol). Membranes were ponceau stained and imaged. Membranes were blocked in 5% milk in Tris-buffered saline with Tween (TBST) for 1 h and then probed with primary antibody overnight (**Table 1**). Membranes were washed with TBST for 30 min, incubated with secondary antibodies conjugated to horseradish peroxidase (α-mouse or α-rabbit; 1:5,000) at room temperature for 1 h, washed with TBST for 30 min, and detected using Clarity Western ECL Substrate (1705061; Bio-Rad) and Chemidoc MP Imaging System (Bio-Rad). Images were formatted using Adobe Photoshop and Illustrator.

**Table 1.** Antibodies used for western blot and immunofluorescence.

| Antibody | Source | Identifier | Use (concentration) |
| --- | --- | --- | --- |
| Mouse anti-CMV IE1/2 | Virusys Corporation | Cat: p1215 | WB (1:5,000) |
| Mouse anti-CMV UL44 | Virusys Corporation | Cat: CA006-100 | WB (1:36,000) |
| Mouse anti-CMV pp28 | Virusys Corporation | Cat: CA004-1 | WB (1:4,000)<br>IF (1:250) |
| Mouse anti-CMV pp65 | Virusys Corporation | Cat: CA003-100 | WB (1:2,000) |
| Mouse anti-CMV gB | Virusys Corporation | Cat: CA005-100 | WB (1:4,000) |
| Rabbit anti-macroH2A1 | Abcam | Cat: 37264 | WB (1:1,000) |
| Mouse anti-Vinculin | Sigma Aldrich | Cat: V9131 | WB (1:10,000) |
| Mouse anti-α-Tubulin | Thermo Fisher Scientific | Cat: 32-2500 | IF (1:1000) |
| Mouse anti-α-Tubulin | Santa Cruz Biotechnology | Cat: sc-69969 | WB (1:1,000) |
| Mouse anti-β-actin | Abcam | Cat: 5441 | WB (1:10,000) |
| Rabbit anti-KIF1A | Abcam | Cat: ab180153 | WB (1:2,500) |
| Rabbit anti-KIBRA (WWC1) | Cell Signaling Technologies | Cat: 8774S | WB (1:1,000) |
| Rabbit anti-H3 | Abcam | Cat: 1791 | WB (1:20,000) |
| Rabbit anti-H3K27me3 | Cell Signaling Technologies | Cat: 9733T | WB (1:100) |
| Peroxidase-AffiniPure goat anti-rabbit | Jackson ImmunoResearch Laboratories | Cat: 111-035-045 | WB (1:5,000) |
| Peroxidase-AffiniPure goat anti-mouse | Jackson ImmunoResearch Laboratories | Cat: 115-035-003 | WB (1:5,000) |
| Goat anti-mouse IgG (H+L) AlexaFluor 555 | Thermo Fisher Scientific | Cat: A-32727 | IF (1:300) |

### Quantification of HSV-1 genomes by droplet digital (ddPCR)

Quantification was carried out as previously described^22^. In brief, cells were harvested at the indicated times after infection by trypsinization, washed with 1× PBS, and centrifuged at 5,000 × *g* for 2 min. Pellets were flash-frozen in liquid nitrogen and stored at −80°C until processed. HCMV DNA within cells was isolated from frozen pellets using QIAamp DNAMini Kit (51304; Qiagen).

Supernatants were harvested at the indicated times after infection, centrifuged at >3,500 × *g,* and filtered through 40-µm sterile syringe filters. DNA on the exterior of filtered capsids was digested for 1 h at 25°C with 20.3 units DNase (79254; Qiagen) supplemented with 10 mM MgCl_2_. DNase was inactivated at 75°C for 10 min followed by vortexing. Capsids were then digested with 3 mg/ml proteinase K (BP1700; Thermo Fisher Scientific) in 100 mM KCl, 25 mM EDTA, 10 mM Tris-HCl pH 7.4, and 1% Igepal for 1 h at 50°C. HCMV genomes from digested capsids were isolated using QIAamp DNAMini Kit.

A duplexed droplet digital PCR was performed to measure the levels of cellular or supernatant HCMV genomes on the QX100 droplet digital PCR system (Bio-Rad Laboratories) using a primer/probe set specific to HCMV UL55. Cell numbers were determined using a primer/probe set specific to human Beta-globin, a reference gene that exists at two copies per cell. The ddPCR reaction mixture consisted of 12.5 µl of a 2× ddPCR Supermix for Probes no dUTP (1863024; Bio-Rad), 1.25 µl of each 20× primer-probe mix (**Table 2**), and 10 µl of template DNA. 20 µl of each reaction mixture was loaded onto a disposable plastic cartridge (1864008; Bio-Rad) with 70 µl of droplet generation oil (1863005; Bio-Rad) and placed in the droplet generator (Bio-Rad). Droplets generated were transferred to a 96-well PCR plate (12001925; Bio-Rad), and PCR amplification was performed on a Bio-Rad C1000 Touch Thermal Cycler with the following conditions: 95°C for 10 min, 40 cycles of 94°C for 30 s, and 60°C for 1 min, followed by 98°C for 10 min, and ending at 4°C. After amplification, the plate was loaded onto the droplet reader (QX200; Bio-Rad) and the droplets from each well of the plate were automatically read with droplet reader oil (186–3004; Bio-Rad) at a rate of 32 wells per hour. Data were analyzed with QuantaSoft analysis software and the quantitation of target molecules presented as copies per microliter of the PCR reaction. HCMV genome values were standardized to cellular β-globin levels. Experiments were completed in biological triplicate and statistical analysis was performed as indicated in figure legends using Prism v10 (GraphPad Software).

**Table 2:**
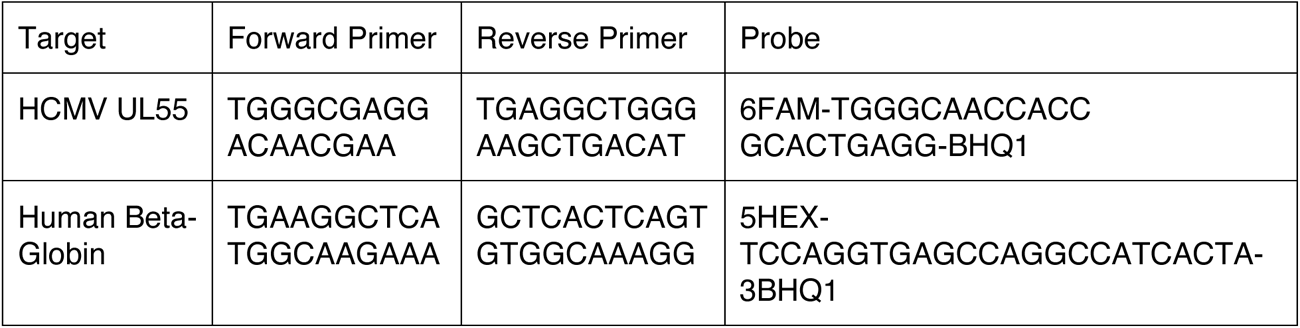
Primers and probes for ddPCR.

### RNA sequencing

Three biological replicates per time point were obtained from independent infections. Cells were harvested at the indicated times after infection by trypsinization, washed with PBS, and centrifuged at 5,000 × *g* for 2 min. RNA was harvested using New England BioLabs Monarch® Total RNA Miniprep Kit (T2010S) as per kit instructions.

RNA was quantified by Nanodrop and integrity was analyzed with the 4200 Tapestation Bioanalyzer system (Agilent). 500 ng of total RNA with an RNA Integrity Number (RIN) >9.5 were used to prepare sequencing libraries with the TruSeq Stranded mRNA Library Prep Kit (20020594; Illumina). Library concentrations were measured with Qubit dsDNA HS Assay Kit (<u>Q32854</u>; Thermo Fisher Scientific) and then analyzed with Agilent High Sensitivity D5000 ScreenTape System and pooled. Libraries were sequenced with 100-bp paired-end reads on an Illumina NextSeq 2000 sequencer at the Fred Hutch Genomics Core Facility.

### RNA-seq analysis

A concatenated fasta file was created using cDNA sequences from release 110 of Ensembl for the human genome and *Towne-HCMV* genome generated from sequencing map in Murphy e*t al.* 2003^59^ and genbank sequences, which was then used to construct a Salmon index^60^. Expression for each transcript was quantified from raw reads using Salmon v1.9 with libType set as automatic. DESeq2^61^ (v1.30.1) in R (v4.0.3) was used first to perform all pairwise comparisons for WT and macroH2A1 KO both across time points and between the two genotypes. There were six time points each in biological triplicates, resulting in 36 datasets: six time points compared against each other for each genotype (15x2) and six time points compared between WT and macroH2A1 KO (6) A superset of genes was made by combining lists of genes with adjusted p-value ≤ 0.05 from each comparison. The expression matrix across genotypes and time points was transformed using the “rlog” function in DESeq2, and then the expression values for the superset of genes were extracted. The normalized read count for each gene was averaged across replicates, Z-transformed across time points and genotypes, and then the matrix of Z-scores was subjected to k-means clustering (k=4). Raw reads for ENCODE datasets were obtained and quantified with Salmon in the same manner as the CMV samples. CMV samples and ENCODE samples were loaded together in DESeq2 as a single DESeq dataset and transformed using the “rlog” function. Normalized expression values for genes in cluster 1 were then extracted to plot the distance matrix and principal components. The distance matrix was calculated using the “dist” function in R and plotted using the pheatmap package. Principal component analysis was performed using the “prcomp” function in R and plotted using ggplot2. GO enrichment analysis was performed with WebGestalt^62^. macroH2A and H3K27me3 enrichment at genes from different clusters were performed using published data for WT and macroH2A.1 KO HFF cells. Enrichment was calculated across the whole gene (Gene start and Gene end definitions from Ensembl).

### Targeted siRNA screen

HFFs were plated in 6-well plates and infected at MOI of 1 with HCMV-GFP Towne as described. Cells were transfected at 12 and 24 hours post infection with 25 pmol/well siRNA (Silencer select, Ambion distributed by Thermo Fisher) (**Table 3**) using lipofectamine RNAiMAX (Thermo Fisher). Non-targeting “Negative Control #1” (Cat. 4390843) was used as siRNA control. At 4 days post infection, supernatants were collected and flash frozen in liquid nitrogen for GFP-foci and plaque assay. Cells were pelleted and split, 75:25 for protein lysate and RNA respectively. Cells for protein lysate were lysed as previously described. Cell pellets for RNA extraction were flash frozen in liquid nitrogen and stored at -80C.

### RT-qPCR

RNA was extracted by TRIzol (Invitrogen) and cDNA was generated using Iscript Reverse Transcription kit (Bio-Rad). RT-qPCR was performed using a CFX384 Touch Real-Time PCR Detection System (Bio-Rad) and iTaq Universal SYBR Green One-Step kit (Bio-Rad). Primers for RT-qPCR are described in Table 3.

**Table 3.**
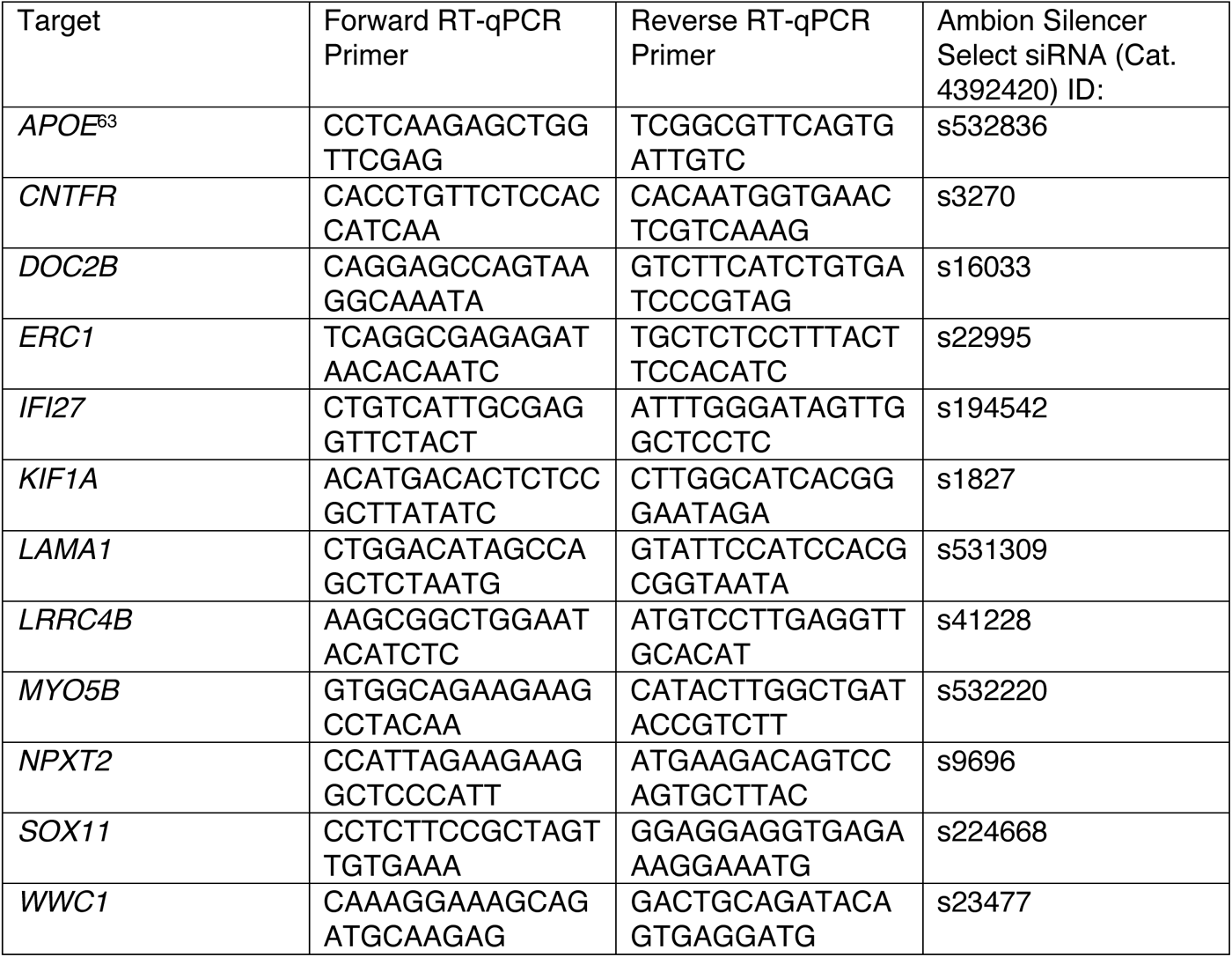
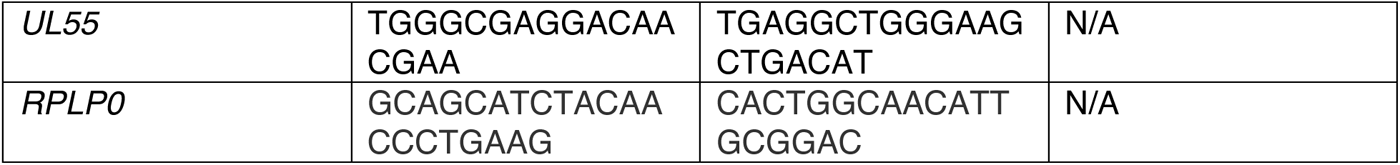
RT-qPCR primers and siRNA IDs.

### Immunofluorescence

As previously described^8^, in brief: Cells were plated on poly-L coated glass coverslips the day prior to infection. Cells were then infected with HCMV at an MOI of 1 and collected at 72 hpi. For harvest, cells were fixed with cold 4% PFA in 1× PBS for 15 min. Cells were permeabilized with 0.5% Triton-X in 1× PBS for 15 min, then blocked in 10% human serum in 1× PBS for 1 h, incubated with primary antibody (diluted as noted in **Table 2**) in 10% human sera in 1× PBS for 1 h. Slides were incubated with secondary antibodies at a dilution of 1:300 in 3% BSA in 1× PBS for 1 h. Coverslips were fixed to microscope slides with Invitrogen ProLong Gold Antifade Mountant or Vectashield (Tubulin stain). Images were taken on Leica Stellaris Confocal with 63× oil objective at room temperature. Images were formatted using Adobe Photoshop and Illustrator.

Images were analyzed using Bitplane Imaris v9.1.1. Labeled nuclei and vIACs were segmented using the Surfaces tool on DAPI and pp28 stains respectively, and average volumes, sphericity and mean fluorescence intensities were reported for each field.

### Plaque area quantification

Once plaque assays were fixed (as per Cells and Virus section), plates were imaged on a Molecular Devices ImageXpress Micro high-content imaging system equipped with a Nikon 4x/0.2 Plan Apo objective. To provide a quantitative measurement of staining intensity, we acquired fluorescence images with a Cy5 filter set. Twenty-four overlapping fields were imaged per well, providing 73% coverage of the total well area.

Using FIJI is Just ImageJ version 2.1.0: Java 1.8.0_172 [64-bit] threshold was set to top 8% of pixel intensity (to account for well-to-well variation in crystal violet staining). The image was converted to binary and “analyze particles” feature was used to acquire area for particles over 1000 pixel units squared (**Sup Figure 6A**).

We also acquired transmitted light images of selected samples on a Nikon Eclipse Ti inverted microscope equipped with a Nikon 4x/0.2 Plan Apo objective. Images were acquired on a Photometrics Prime BSI Express sCMOS camera through an orange (580-610 nm) filter. This wavelength range was chosen to match the absorbance spectrum of crystal violet, thus maximizing staining contrast and ensuring high dynamic range.

### Electron microscopy

Cells were fixed in 2% paraformaldehyde and 2.5% glutaraldehyde in 0.1 M sodium cacodylate buffer (pH 7.3) at 4°C. Fixed cells were rinsed briefly in 1% sucrose in 50 mM cacodylate (pH 7.2), then postfixed on ice for 30 min in a solution of 1% osmium tetroxide (RT19152; EM Sciences) and 0.8% potassium ferricyanide in 50 mM cacodylate (pH 7.2). Cell pellets were washed twice briefly at 25°C in 1% sucrose in 50 mM cacodylate (pH 7.2) and then washed in three changes of 50 mM cacodylate (pH 7.2) for 5 min each. Cell pellets were treated with 0.2% tannic acid (1401-55-4; Sigma-Aldrich) in 50 mM cacodylate (pH 7.2) for 15 min at 25°C and then rinsed several times in water. Cells were dehydrated through a graded ethanol series and embedded in Epon 12 resin (18010; Ted Pella). 70-nm thin sections were cut using an Ultracut UC7 ultramicrotome (Leica Mikrosysteme) and collected on 200 mesh formvar/carbon coated copper grids (01800; Ted Pella). Sections were stained with 2% aqueous uranyl acetate and Reynolds lead citrate. Cell pellet sections were imaged using a Talos L120C microscope operated at 120 kV with a Ceta-16 M (4,096 × 4,096) camera (Thermo Fisher Scientific).

All data were collected at spot size 5 with a 100-µm C2 aperture and 70-µm objective aperture. Images were formatted using Adobe Photoshop and Illustrator.

Image analysis was done using FIJI is Just ImageJ version 2.1.0: Java 1.8.0_172 [64-bit]. For analysis 2.5 micron by 2.5 micron grids were drawn on non-overlapping regions of cytosol. “Freehand line tool” was used to trace all lengths of ER in the field of view and length of line was measured with ImageJ “Measure” function (**Sup Figure 2B**). Macro code for vIAC subcompartments can be found at: **10.5281/zenodo.11521560** (**Sup Figure 2D**).

### Statistical Analysis

Graphs were generated and statistical analysis were run as marked in figure legends using GraphPad Prism v 9.1.2.

